# Evaluating the burstlet theory of inspiratory rhythm and pattern generation

**DOI:** 10.1101/718759

**Authors:** Prajkta S. Kallurkar, Cameron Grover, Maria Cristina D. Picardo, Christopher A. Del Negro

**Affiliations:** Department of Applied Science, Integrated Science Center, 540 Landrum Dr., William & Mary, Williamsburg, VA 23185

## Abstract

The preBötzinger complex (preBötC) generates the rhythm and rudimentary motor pattern for inspiratory breathing movements. Here, we test ‘burstlet’ theory (Kam, Worrell, Janczewski, et al. 2013), which posits that low amplitude burstlets, subthreshold from the standpoint of inspiratory bursts, reflect the fundamental oscillator of the preBötC. In turn, a discrete suprathreshold process transforms burstlets into full amplitude inspiratory bursts that drive motor output, measurable via hypoglossal nerve (XII) discharge *in vitro*. We recap observations by Kam & Feldman: field recordings from preBötC demonstrate bursts and concurrent XII motor output intermingled with lower amplitude burstlets that do not produce XII motor output. Manipulations of excitability affect the relative prevalence of bursts and burstlets and modulate their frequency. Whole-cell and photonic recordings of preBötC neurons suggest that burstlets involve inconstant subsets of rhythmogenic interneurons. We conclude that discrete rhythm- and pattern-generating mechanisms coexist in the preBötC and that burstlets reflect its fundamental rhythmogenic nature.

## INTRODUCTION

Breathing is a vital rhythmic behavior. Inspiration, the inexorable active phase of the breathing cycle, originates from the preBötzinger complex (preBötC) in the lower medulla (Del Negro, Funk, and Feldman 2018; Jeffrey C Smith et al. 1991). preBötC interneurons generate rhythmic activity and project to cranial and spinal premotor and motor neurons that drive inspiratory muscles. However, the neural mechanisms that give rise to inspiratory rhythm and motor pattern are unclear. These mechanisms can be studied in reduced preparations that isolate the preBötC and cranial hypoglossal (XII) inspiratory motor circuits, and thus provide an experimentally advantageous minimal breathing-related microcircuit *in vitro*. Here we disentangle neural mechanisms intrinsic to the preBötC that engender both inspiratory rhythm and fundamental aspects of the motor pattern.

We stipulate: preBötC bursts propel inspiratory activity to premotor and motor neurons that drive motor output thus they are important for motor pattern. However, are preBötC bursts rhythmogenic? The following observations cast doubt on that premise. The magnitude of inspiratory bursts (in the preBötC and motor output) can be diminished while only minimally affecting their frequency (Del Negro et al. 2002; Shereé M. Johnson, Koshiya, and Smith 2001; Pace et al. 2007; Peña et al. 2004). Also, manipulating excitability in the preBötC affects the frequency, but not the magnitude of preBötC inspiratory bursts and motor output (Del Negro et al. 2009; Sun et al. 2019). It thus appears that two discrete phenomena emanate from the preBötC: a fundamental rhythm (whose frequency is adjustable) and a rudimentary pattern consisting of bursts that drive motor output. These appear to be separable processes as codified by Kam and Feldman in their *burstlet* hypothesis (Feldman and Kam 2015; Kam, Worrell, Janczewski, et al. 2013). To describe one cycle, preBötC neurons experience a ramp-like depolarization lasting 100-400 ms, which reflects recurrent excitatory synaptic activity among constituent rhythmogenic neurons (Kam, Worrell, Ventalon, et al. 2013; J. C. Rekling, Champagnat, and Denavit-Saubie 1996; J. C. Smith et al. 1990). This is called the preinspiratory phase because it precedes – and ordinarily leads to – the inspiratory burst. However, Kam and Feldman showed that preinspiratory activity can be divorced from inspiratory bursts by lowering the neural excitability. What often remains is preBötC network activity matching that of the preinspiratory phase but absent the burst; they dubbed these events *burstlets* (Kam, Worrell, Janczewski, et al. 2013).

Here, we test the burstlet hypothesis of inspiratory rhythm and pattern generation by explicitly or conceptually repeating Kam and Feldman’s (2013) experiments. We, too, detected preBötC burstlets absent XII output, which were distinct from larger amplitude preBötC bursts accompanied by XII output. Manipulations that lower neural excitability increase the prevalence of burstlets relative to bursts. Composite preBötC rhythm was voltage dependent because manipulations of excitability control its frequency. Intracellular recordings and photonic imaging of preBötC inspiratory neurons demonstrate that the burstlets occur in subsets of preBötC neurons, not the entire rhythmogenic population that participates in bursts. Our results also support the fundamental tenet of burstlet theory that preinspiratory activity and burstlets reflect a common rhythmogenic mechanism, and that a threshold process causes burstlets (i.e., preinspiratory activity) to trigger bursts and subsequent motor output. These results imply that pattern generation, although a distinct process from rhythm generation, starts from the preBötC core microcircuit.

## MATERIALS AND METHODS

The Institutional Animal Care and Use Committee at William & Mary approved these protocols, which conform to the policies of the Office of Laboratory Animal Welfare (National Institutes of Health, Bethesda, MD) and the guidelines of the National Research Council (National Research Council 2011). Mice were housed in colony cages on a 14-hour light/10-hour dark cycle with controlled humidity and temperature at 23° C and were fed *ad libitum* on a standard commercial mouse diet (Teklad Global Diets, Envigo, Madison, WI) with free access to water.

### Mice

preBötC field recording experiments employed CD-1 mice (Charles River, Willmington, MA). Whole-cell recordings employed CD-1 mice as well as mice with Cre-dependent expression of fluorescent Ca^2+^ indicator GCaMP6f dubbed Ai148 by the Allen Institute (RRID:IMSR_JAX:030328) (Daigle et al. 2018). Imaging experiments employed Ai148 mice. We crossed homozygous *Dbx1*^*Cre*^ (Bielle et al. 2005) females with Ai148 males. We refer to their offspring as Dbx1;Ai148 mice. Newborn Dbx1;Ai148 pups express GCaMP6f in neurons derived from progenitors that express the embryonic transcription factor *developing brain homeobox 1* (i.e., *Dbx1*).

### Slice preparations

Neonatal mice (P0-4) of both sexes were anesthetized by hypothermia and killed by thoracic transection. Brainstems were removed in cold artificial cerebrospinal fluid (ACSF) containing (in mM): 124 NaCl, 3 KCl, 1.5 CaCl_2_, 1 MgSO_4_, 25 NaHCO_3_, 0.5 NaH_2_PO_4_, and 30 dextrose, which we aerated with 95% O_2_ and 5% CO_2_. Brainstems were then glued to an agar block with the rostral side up. We cut a single 450-500 µm-thick transverse medullary slice with preBötC at the rostral surface. The position of the preBötC was benchmarked according to neonatal mouse preBötC atlases (Ruangkittisakul, Panaitescu, and Ballanyi 2011; Ruangkittisakul et al. 2014).

### Electrophysiology

Slices were held in place and perfused with ACSF (∼28° C) at 2-4 ml-min^-1^ in a recording chamber on a fixed-stage upright microscope. The external K^+^ concentration, i.e., [K^+^]_o_, in the ACSF was initially raised to 9 mM, which facilitates robust rhythm and motor output in slices (Funk and Greer 2013).

Population activity from preBötC interneurons and XII motor neurons was recorded using suction electrodes fabricated from borosilicate glass pipettes (OD: 1.2 mm, ID: 0.68 mm). preBötC field recordings were obtained by placing the suction electrode over the rostral face of the preBötC at the surface of the slice. XII motor output was recorded from XII nerve rootlets, which are retained in slices. Signals were amplified by 20,000, band pass filtered at 0.3-1 kHz, and then RMS smoothed using a differential amplifier (Dagan Instruments, Minneapolis, MN). Smoothed signals were used for display and quantitative analyses.

We used an EPC-10 patch-clamp amplifier (HEKA Instruments, Holliston, MA) for whole-cell current-clamp recordings. Patch pipettes were fabricated from borosilicate glass (OD: 1.5 mm, ID: 0.86 mm) to have tip resistance of 4-6 MΩ. The patch pipette solution contained (in mM): 140 K-gluconate, 5 NaCl, 0.1 EGTA, 10 HEPES, 2 Mg-ATP, 0.3 Na_3_-GTP. We added 50 µM Alexa 488 hydrazide dye (A10436, Life Technologies, Carlsbad, CA) for visualization after whole-cell dialysis. Whole-cell recordings were made from preBötC inspiratory neurons selected visually based on rhythmic fluorescence changes in Dbx1;Ai148 mice. In CD-1 mice, we only collected whole-cell data from preBötC inspiratory neurons that were active in sync with XII motor output. Membrane potential trajectories were low-pass filtered at 1 kHz and digitally recorded at 4 kHz using PowerLab data acquisition system, which includes a 16-bit analog-to-digital converter and LabChart v7 software (ADInstruments, Colorado Springs, CO).

We modified [K^+^]_o_ in the ACSF from 9 to 3 mM to modulate the excitability in the preBötC. Each trial consisted of a sequence of non-contiguous [K^+^]_o_ levels selected randomly in descending order.

Low-amplitude activity in preBötC field recordings was classified as a burstlet if it met these two criteria: the peak of preBötC activity exceeded the mean of the distribution of baseline noise by two standard deviations (SD, i.e., 2*SD) and there was negligible concurrent activity in the XII root recording. The mean and SD of baseline noise were computed by sampling every data point during a sliding 120 s window, then constructing a histogram of baseline noise, and fitting the mean and SD from that distribution.

For field recordings, we measured the amplitude and area of the preBötC population activity and XII motor output, as well as their frequencies using LabChart peak parameters plugin (ADInstruments). For whole-cell recordings, inspiratory bursts refer to depolarizations with concomitant spiking in preBötC neurons that occur in sync with XII motor output. We measure the frequencies of preBötC neuronal activity and XII motor output using LabChart plugins (ADInstruments). To measure their amplitude and area, inspiratory bursts were digitally smoothed using smoothing function in LabChart (ADInstruments) to minimize the spikes but preserve the underlying envelope of depolarization, which we define as the inspiratory drive potential (Pace et al. 2007).

We wrote algorithms in MATLAB (MathWorks, Natick, MA) to calculate the mean frequency (f) (or cycle time) as well as the amplitude (amp) and area of bursts and burstlets. The coefficient of variation (CV) of preBötC or XII motor output frequency was calculated as the ratio of standard deviation to the mean frequency.

Cycle-triggered averages were calculated and plotted in IgorPro (v.8, Wavemetrics, Oswego, OR) using the onset of XII output as the event trigger for averaging preBötC inspiratory bursts; the onset of the burstlet itself served as the event trigger for averaging burstlets. We obtained the depolarization rate (V/s) of the event-triggered averages as the quotient of event amplitude and the elapsed time for that event to reach its peak amplitude. For preinspiratory activity, event amplitude was calculated as the absolute difference between baseline at the onset burst and field amplitude at the onset of XII motor output. For burst, event amplitude was calculated as the difference between peak amplitude and field amplitude at the onset of XII motor output. For burstlet, event amplitude was calculated as the difference between peak amplitude and baseline at the onset of the burstlet.

### Two-photon Imaging

We imaged intracellular Ca^2+^ in neurons contained in slices from Dbx1;Ai148 mice using a multi-photon laser-scanning microscope (Thorlabs, Newton, NJ) equipped with a water immersion 20x, 1.0 numerical aperture objective. Illumination was provided by an ultrafast tunable laser with a power output of 1050 mW at 970 nm, 80 MHz pulse frequency, and 100 fs pulse duration (Coherent Chameleon Discovery, Santa Clara, CA). We scanned Dbx1;Ai148 mouse slices over the preBötC and collected time series images at 32 Hz. Each frame reflects one-way raster scans with a resolution of 256 × 256 pixels (116 × 116 µm). Fluorescence data were collected using Thorlabs LS software and then analyzed using Fiji (Schindelin et al. 2012; Schneider, Rasband, and Eliceiri 2012), MATLAB, and IgorPro.

Regions of interest (ROIs) were detected using MATLAB. First we found the individual cycle periods between network bursts in each Dbx1;Ai148 slice by averaging the fluorescence intensity of all pixels for each frame, and then calculating the time between the fluorescence peaks. The collection of cycle periods is normally distributed; the 96% confidence intervals are defined by the mean cycle period ±2*SD.

Next, we down-sampled the planar resolution of our stack of images by 2^n^ (n ≥ 1). The mean fluorescence of the constituent pixels was assigned to each composite pixel. We performed temporal fast Fourier transforms (FFTs) on the composite pixels. The maximum FFT value within the 96% confidence intervals for slice frequency (determined above) was then mapped to a corresponding position in a new processed 2D image. This method quantifies how strongly a composite pixel changes fluorescence at frequencies that correspond to inspiratory rhythm. After having created the complete processed image, we computed the mean and SD for FFT values associated with all composite pixels. Any composite pixel with intensity less than mean + 2*SD was set to zero. Any contiguous remaining pixel sets (whose diameter exceeds 6 µm) were retained as ROIs.

We then applied the set of ROIs to analyze Ca^2+^ transients in the original fluorescence imaging stack using the equation (F_i_ – F_0_)/F_0_, i.e., ΔF/F_0_, where F_i_ is the instantaneous average fluorescence intensity of all the pixels in a given ROI and F_0_ is the average fluorescence intensity of all the pixels within the same ROI averaged over the entire time series. Finally, we smoothed the ΔF/F_0_ time series with a forward moving average with a window of four time points.

### Statistics

All the statistical hypothesis tests were calculated using Prism8 (GraphPad, San Diego, CA). Changes in the frequency and amplitude of preBötC field activity and XII output as a function of [K^+^]_o_ were evaluated using linear regression. We compared group means using either Student’s paired t test or repeated measures one-way ANOVA, applying Holm-Sidak’s multiple comparison test post-hoc. We compared the variability of frequency of preBötC events and XII motor output using a Kruskal-Wallis test, applying Dunn’s multiple comparison test post-hoc. We compared the frequency of preBötC events using Welch’s t test.

## RESULTS

### preBötC generates bursts at high levels of excitability but burstlets appear as excitability decreases

We manipulated excitability by varying [K^+^]_o_ by integer units between 9 and 3 mM. At high [K^+^]_o_ (e.g., 9 or 7 mM in Fig. 1A) we observed mostly burst events, defined by peaks of activity in the preBötC field recording with coincident XII output. Nevertheless, we also observed events in the preBötC field recording whose amplitudes measured 15-65% of the bursts and occurred without coincident XII discharge (Fig. 1A).

**Figure 1.**
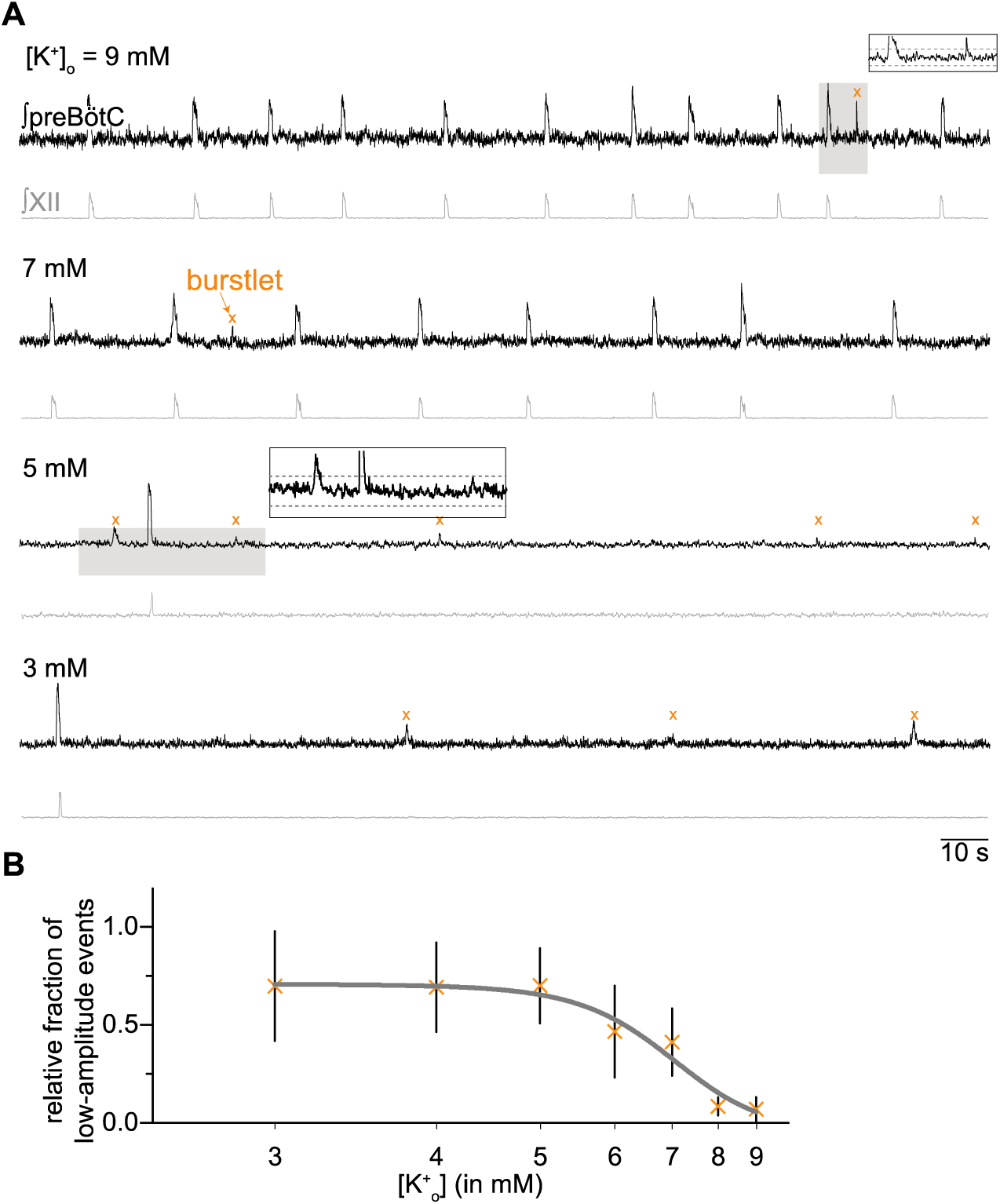
preBötC field and XII recordings demonstrate inspiratory burst and ‘burstlet’ rhythms. A, preBötC field (black) and XII (gray) at different levels of [K^+^]_o_. Insets show expanded traces from gray boxes. Dashed lines in the insets mark the 95% CI. Orange X’s symbols indicate burstlets. Time calibration applies to all traces. B, The relative fraction burstlets in preBötC field recordings as a function of [K^+^]_o_.

We detected and then studied low-amplitude preBötC events whose peaks exceeded the 95% CIs of baseline noise (Fig. 1A, insets). That detection process ensures that low-amplitude events are unlikely (with probability < 0.05) to be ordinary uncoordinated fluctuations of neural activity. The alternative is that these low-amplitude preBötC events reflect coherent network activity.

If the low-amplitude events reflect burstlets as defined by Kam and Feldman (Kam, Worrell, Janczewski, et al. 2013), then they should be more abundant at low levels of excitability where the collective activity of rhythmogenic neurons may not reach the threshold for burst generation. Visual inspection of the traces in Fig. 1A show that to be the case, i.e., low-amplitude events devoid of XII output are more abundant at 5 and 3 mM [K^+^]_o_ compared to 9 and 7 mM [K^+^]_o_.

At 3 mM [K^+^]_o_, 70% ± 3% (n = 12 slices) of detected preBötC events occurred without concomitant XII output. We quantified the relative abundance of low-amplitude vs. burst events for the entire data set (Fig. 1B, n = 19 slices). At incrementally higher [K^+^]_o_ levels, the relative fraction of low-amplitude events decreases in a sigmoidal fashion such that they comprise only 5.2% ± 6.3% (n = 19 slices) of the preBötC events at 9 mM. We conclude that low-amplitude preBötC events, absent motor output, reflect burstlets as defined previously (Kam, Worrell, Janczewski, et al. 2013).

### Voltage dependence of burst-burstlet rhythm

We measured the frequency of preBötC rhythm (f_preBötC_, which we refer to as composite rhythm because the measured events consist of either bursts or burstlets) at different [K^+^]_o_ levels (Fig. 2A) similar to Kam *et al.* (2013a). They measured the frequency of preBötC composite rhythm at three discrete [K^+^]_o_ levels (3, 6, and 9 mM) and reported that frequency was lower at 3 compared to either 6 or 9 mM, yet there was no difference between the frequency measured at 6 vs. 9 mM (see Table 1 of Kam et al., 2013a).

**Table 1.** Effects of [K+]_o_ on the frequency and amplitude of preBötC events and XII motor output. Change in frequency and amplitude of preBötC events and XII output as [K^+^]_o_ is changed from 3 to 9 mM, in 1 mM steps. All results were analyzed using linear regression analysis. Number of slices (n) at each [K^+^]_o_ (mM,n): (3,12); (4,9); (5,8); (6,12); (7,8); (8,8); (9,19).

|  | Slope<br>(Hz/mM or<br>mV/mM) | p-value | R<sup>2</sup> | Figure |
| --- | --- | --- | --- | --- |
| preBötC composite frequency (Hz) | 0.021 | 7.0E-15 | 0.55 | 2A |
| Burst amplitude (mV) | 0.6 | 0.62 | 0.24 | 3A |
| Burstlet amplitude (mV) | 1.6 | 0.001 | 0.15 | 3B |
| XII amplitude (mV) | 1.5 | 0.50 | 0.47 | 3C |

**Figure 2.**
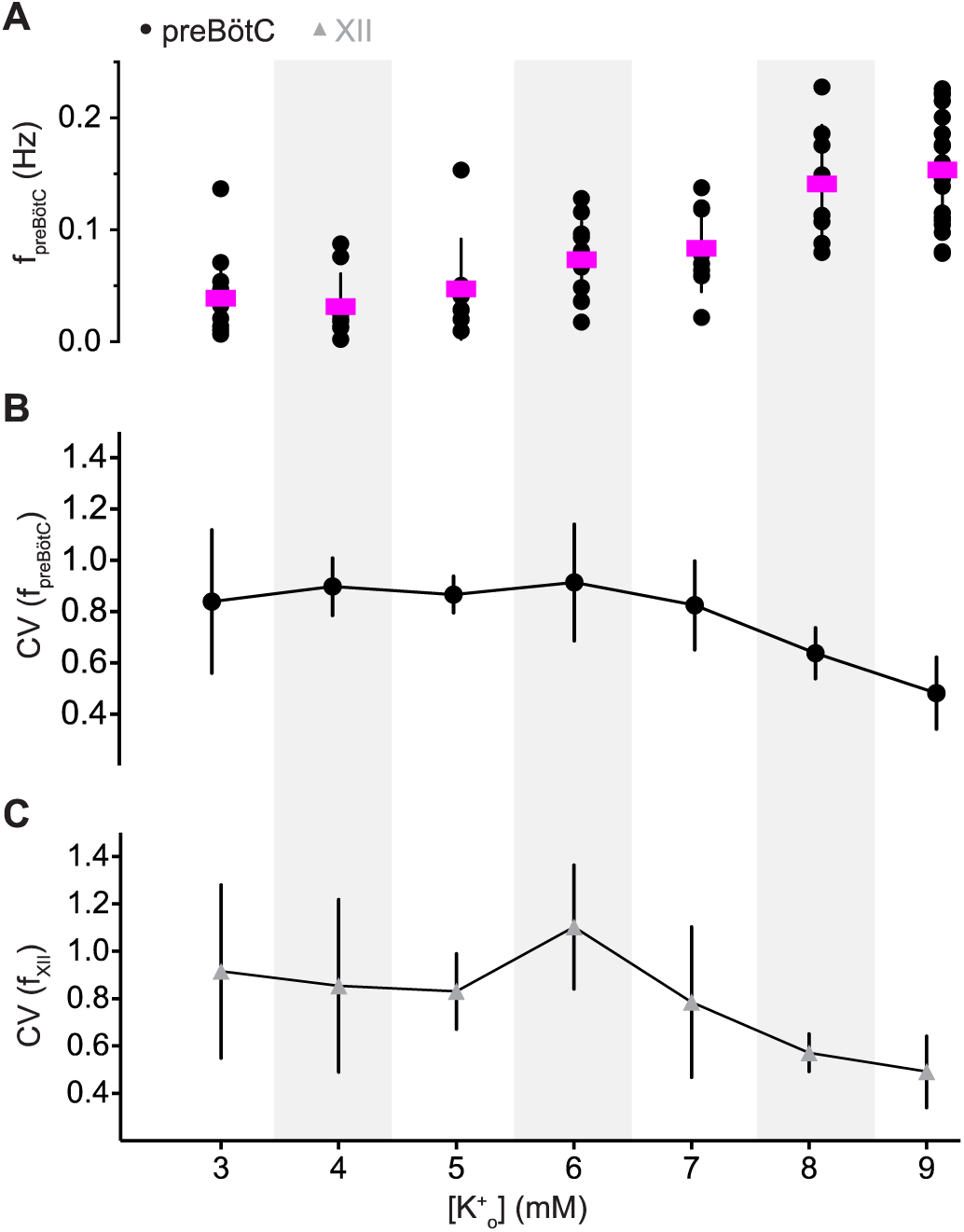
Frequency and its variability of preBötC composite rhythm and XII motor output. A, frequency of composite (f_preBötC_, filled circles) rhythm as a function of external K^+^ concentration, i.e., [K^+^]_o_. The frequency from each slice preparation is shown with mean frequency (magenta) and SD (vertical lines). B-C, mean coefficient of variation (CV) for composite (filled circles) and XII (gray triangles) rhythms as a function of [K^+^]_o_. Vertical bars show SD. Light gray background shading was applied to differentiate [K^+^]_o_ levels.

Those data do not resolve whether the composite rhythm is voltage dependent so we measured rhythmic activity at all integer [K^+^]_o_ levels between 3 and 9 mM. The frequency of the preBötC composite rhythm increased linearly as the excitability increased (Fig. 2A and Table 1, which reports linear regression tests). These data show that preBötC composite rhythm is voltage dependent.

Changes in the excitability of the preBötC affect the periodic variability of XII motor output (Del Negro et al. 2009). We reexamined that principle and further tested variability of composite preBötC rhythm (Fig. 2B). The variability in the frequency of XII motor output, quantified by CV, peaked at ∼1.1 when [K^+^]_o_ was 6 mM (Fig. 2C and Table 2). In contrast, the CV of the frequency of the preBötC composite rhythm remained between 0.9-0.7 over low to medium levels of excitability (3-6 mM [K^+^]_o_) without peaking at 6 mM [K^+^]_o_ (Fig. 2B, Table 2). These data suggest that peak CV in the XII output frequency at 6 mM [K^+^]_o_, an intermediate level of excitability, is not attributable to instability in the preBötC rhythm. It rather reflects the equal probability of evoking either burstlets (absent XII output) and bursts with XII output (see our Fig. 1B or Fig. 2 of Kam et al., 2013), which makes the periodic XII output more variable than preBötC activity.

**Table 2.** Effects of [K+]_o_ on the variability of frequency of preBötC composite rhythm and XII motor output. The reported p-values (calculated using Kruskal Wallis test) compare frequency variability of composite rhythm and XII motor output as [K^+^]_o_ is changed from 3 to 9 mM. Post-hoc analysis (calculated using Dunn’s test) report p-values for the group that were statistically significant.

|  | p-value | post-hoc analysis | Figure |
| --- | --- | --- | --- |
| CV (preBötC composite frequency) | 0.003 | 6 vs 9 mM $[K^+]_o$: 0.005 | 2A |
| CV (XII motor output frequency) | 1.408E-5 | 6 vs 8 mM $[K^+]_o$: 0.039<br>6 vs 9 mM $[K^+]_o$: 7.968E-7 | 2B |

Variability is lowest for preBötC composite rhythm, as well as XII output, at 9 mM [K^+^]_o_ (Fig. 2B, Table 2). ACSF containing 9 mM [K^+^]_o_ represents the empirically determined ideal conditions for rhythmically active slices (Funk and Greer 2013; Jeffrey C Smith et al. 1991) where the likelihood of burstlets is minimal (see our Fig. 1B and Table 1 of Kam et al., 2013).

Next, we examined the amplitude of bursts, burstlets, and XII output. The amplitude of preBötC bursts and XII output were invariable over all [K^+^]_o_ levels (Fig. 3A and C), confirmed using linear regression tests (Table 1). However, the burstlet amplitude increased from 1.2 ± 0.6 mV at 3 mM [K^+^]_o_ (n = 12 slices) to 2.3 ± 1.0 mV (n = 13 slices) at 9 mM K^+^ (Fig. 3B), which linear regression showed was unlikely to occur by chance if the slope of burstlet amplitude vs. [K^+^]_o_ was actually zero (p = 0.001, Table 1). These results show that burstlet amplitude is voltage dependent whereas preBötC burst and XII motor output amplitudes are not voltage dependent.

**Figure 3.**
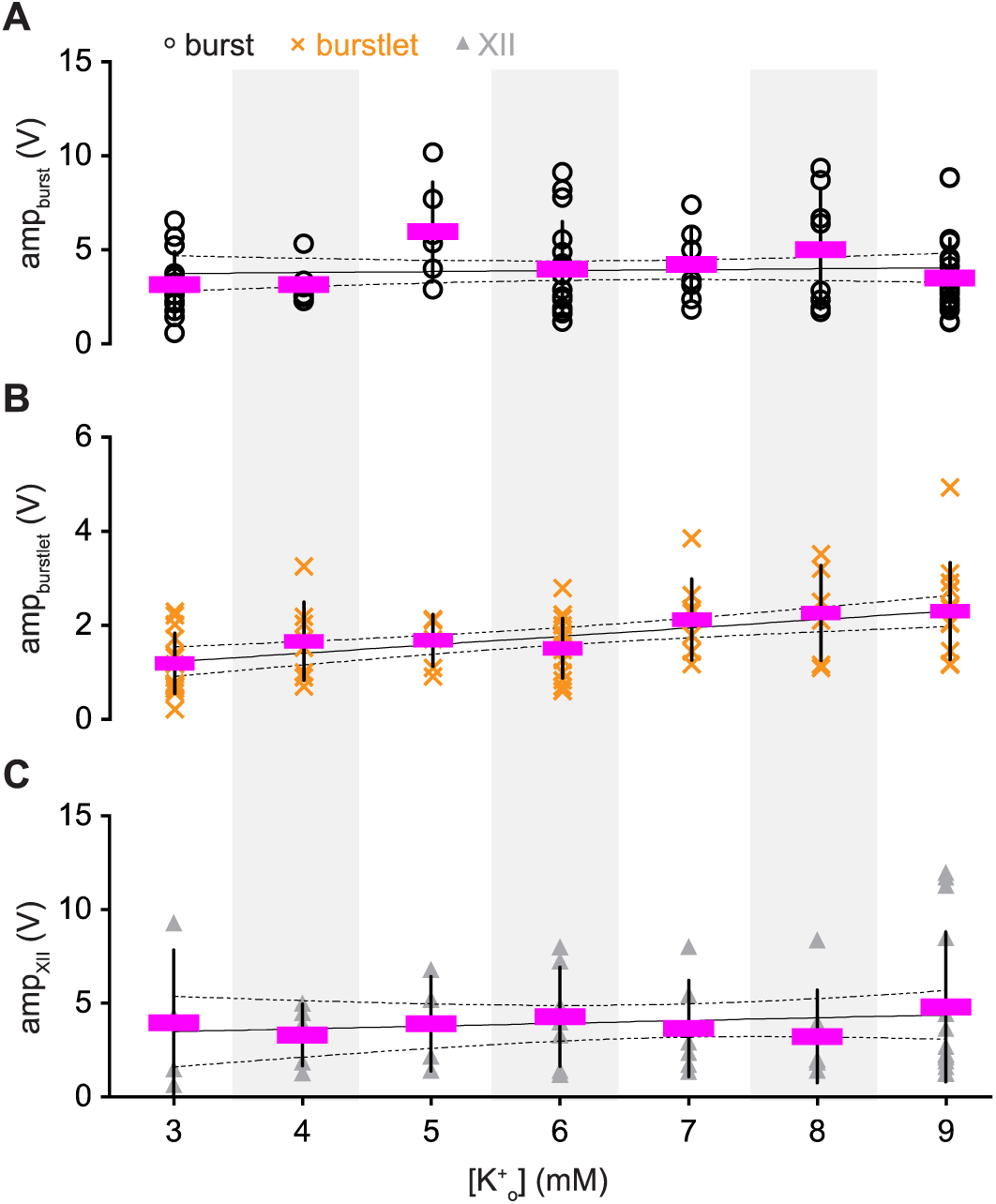
Amplitude of preBötC activity and motor output. A-C, Amplitude of inspiratory bursts (amp_burst_, open circles), burstlets (amp_burstlet_, orange X symbols), and XII output (amp_XII_, gray triangles) as a function of external K^+^ concentration, i.e., [K^+^]_o_. The amplitude of each slice preparation is shown with mean amplitude (magenta) and SD (vertical lines). Linear regression lines (solid) and 95% confidence intervals (dashed) are shown. Light gray background shading was applied to differentiate [K^+^]_o_ levels.

### Burstlets evolve bilaterally and underlie the pre-inspiratory phase of preBötC bursts

If burstlets reflect coherent preBötC rhythmicity, then they should be bilaterally synchronous. To test that prediction, we recorded preBötC activity from both sides of slices along with XII motor output (Fig. 4A). 97% ± 4% and 92% ± 7% of preBötC burstlets were bilaterally synchronous at 6 and 3 mM [K^+^]_o_, respectively (n = 4 slices). The bilateral preBötC bursts commence ∼400 ms before the onset of XII motor output (Fig. 4A inset), which is considered the preinspiratory phase and the hallmark of rhythm generation (Del Negro, Funk, and Feldman 2018; Feldman and Kam 2015; Kam, Worrell, Janczewski, et al. 2013; J. C. Rekling, Champagnat, and Denavit-Saubie 1996; J. C. Smith et al. 1990).

**Figure 4.**
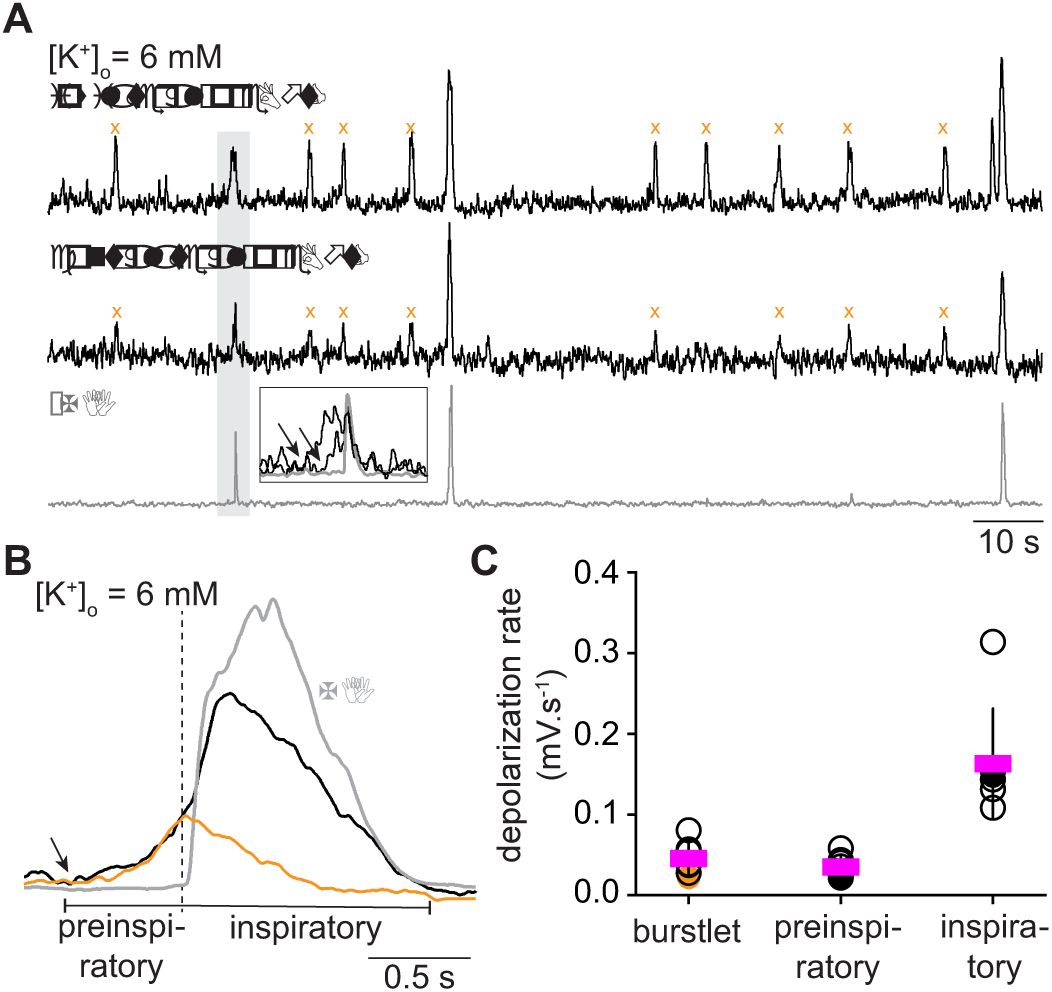
Burstlets are bilaterally synchronous and makeup the preinspiratory phase of bursts. A, bilateral field recordings of the preBötC, with XII output (gray). B, cycle-triggered averages of burstlets (orange), inspiratory bursts (black), and XII output (gray). The onset of preinspiratory preBötC activity and burstlets are marked by an arrow; their peaks occur at the onset of XII motor output, marked by vertical dashed line. For inspiratory bursts, the onset of XII motor output marks the onset of the event. C, plot of rising slope of burstlets, the preinspiratory phase of bursts, and the inspiratory burst itself. The depolarization rate of the rising phase for each slice is shown with individual symbols (filled symbols are shown in the example in B); the mean rising phase is shown in magenta.

If the burstlets reflect the preinspiratory component of the inspiratory burst, as proposed by Kam and Feldman, then their trajectories should look alike when superimposed. At 6 mM [K^+^]_o_, the rising phase of the burstlet resembles the preinspiratory phase of the inspiratory burst (Fig. 4B). We compared the depolarization rates of the rising phase of burstlets, the preinspiratory phase of inspiratory bursts, and the rising phase of inspiratory bursts. The rising phase of burstlets is comparable to the rising phase of the preinspiratory activity, but the rising phase of both burstlets and preinspiratory activity are incommensurate with the rising phase of inspiratory bursts (RM ANOVA, F_(2,12)_ = 33.76, p = 1.2E-5; burstlet vs. preinspiratory, p = 0.577; burstlet vs. inspiratory, p = 2.5E-5; and preinspiratory vs. inspiratory, p = E-5; n = 7 slices) (Fig. 4C). These data suggest that burstlets and the preinspiratory phase of bursts reflect the same underlying process, which is distinct from the process underlying full inspiratory bursts.

### Burstlets are the summation of EPSPs in preBötC neurons

We examined how individual preBötC inspiratory neurons contribute to collective events detected in field recordings (bursts and burstlets) via whole-cell recordings in CD-1 (n = 7) and Dbx1;Ai148 mouse slices (n = 3). Dbx1;Ai148 pups express genetically encoded Ca^2+^ reporter GCaMP6f in *Dbx1*-derived preBötC neurons obligatory for breathing rhythmogenesis (Baertsch, Baertsch, and Ramirez 2018; Bouvier et al. 2010; Cui et al. 2016, 2016; Gray et al. 2010; Vann et al. 2016, 2018; Wang et al. 2014).

In control conditions (9 mM [K^+^]_o_), inspiratory drive potentials synchronized with XII motor output during almost all cycles (96% ± 7%, n = 16 preBötC neurons recorded in slices from 10 different animals). We considered those events bursts. We then modified the excitability by changing [K^+^]_o_ to either 6 mM (n = 11 neurons in 8 slices) or 7 mM (n = 6 neurons in 3 slices). We recorded inspiratory drive potentials of 6-10 mV amplitude that were not accompanied by XII motor output. We considered those events burstlets (Fig. 5A).

**Figure 5.**
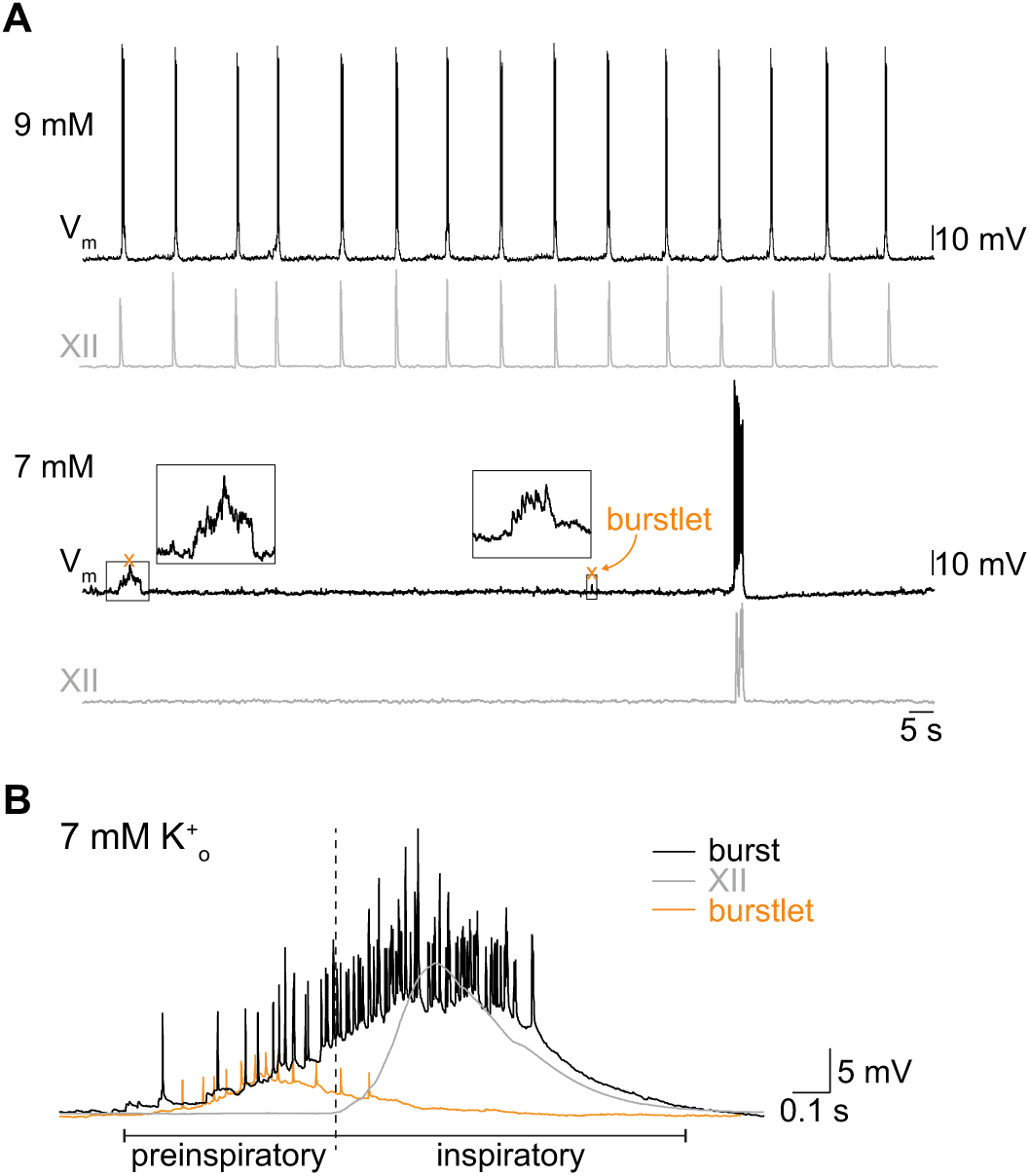
Properties of burstlets recorded intracellularly. A, whole-cell recording (V_m_, top traces) from an inspiratory preBötC neuron with XII output (gray, lower traces) at 9 and 7 mM external K^+^ concentration, i.e., [K^+^]_o_. Insets amplify burstlets at 7 mM [K^+^]_o_. B, cycle-triggered averages of burstlets (orange) and inspiratory bursts (black) recorded intracellularly with XII output (gray). Preinspiratory and inspiratory burst phases are marked. A and B have separate voltage and time calibrations.

During bursts, preBötC neurons, from a baseline membrane potential of −60 mV (below the activation threshold of persistent Na^+^ current, maintained via bias current), exhibit inspiratory drive potentials exceeding 20 mV and intraburst spiking of 2-17 spikes/burst (∼6-60 Hz). During burstlets, these same neurons exhibit excitatory postsynaptic potentials (EPSPs) that summate during the burstlets (Fig. 5A insets) as well as spikes (Fig. 5B shows a cycle-triggered average from a whole-cell recording). We never observed a preBötC neuron (n = 0/17 neurons) that was active during burstlets but not bursts.

Burstlets resemble the preinspiratory phase of bursts. This applies to field recordings (Fig. 4B) and whole-cell recordings at both [K^+^]_o_ levels: 7 mM (n = 4 out of 6 neurons) and 6 mM (n = 6 out of 11 neurons) (Fig. 5B). preBötC field recordings and extracellular unit recordings were shown previously (Ashhad and Feldman 2019; Kam, Worrell, Janczewski, et al. 2013), but these are the first intracellular recordings to show that burstlets reflect the temporal summation of EPSPs, often crossing threshold to generate repetitive spiking, in preBötC neurons.

Kam *et al*., 2013 showed that 89% of the inspiratory preBötC neurons take part in burstlets. We retested that notion by comparing the frequencies of preBötC rhythms monitored during separate whole-cell and field recordings (Fig. 6). We predicted that if 89% of inspiratory neurons participate in burstlets, then the frequency of composite rhythm obtained in whole-cell recordings should be comparable to that obtained in field recordings. We found no difference in the frequencies of preBötC composite rhythms between whole-cell and field recordings at 7 or at 9 mM [K^+^]_o_. However, at 6 mM [K^+^]_o_, f_preBötC_ was significantly lower in whole-cell recordings compared to field recordings (Fig. 6, and Table 3). This difference implies relatively fewer inspiratory preBötC neurons are burstlet-active at 6 mM [K^+^]_o_ compared to 7 or 9 mM.

**Table 3.** Comparison of frequencies of preBötC composite rhythm measured in whole-cell recordings and field recordings at 6, 7 and 9 mM [K+]_o_. Frequencies are reported as mean ± SD. The reported p-values (calculated using Welch’s t test) compare samples of composite frequency measured from the preBötC in either whole-cell or field recordings.

| $[K^+]_o$ (mM) | Composite rhythm | |
| --- | --- | --- |
|  | whole-cell (Hz) | field (Hz) |
|  | p-value |  |
| 6 | 0.04 ± 0.03 | 0.08 ± 0.03 |
|  | 0.04 |  |
| 7 | $0.06 \pm 0.02$ | $0.09 \pm 0.03$ |
|  | 0.21 |  |
| 9 | $0.14 \pm 0.04$ | $0.14 \pm 0.06$ |
|  | 0.65 |  |

**Figure 6.**
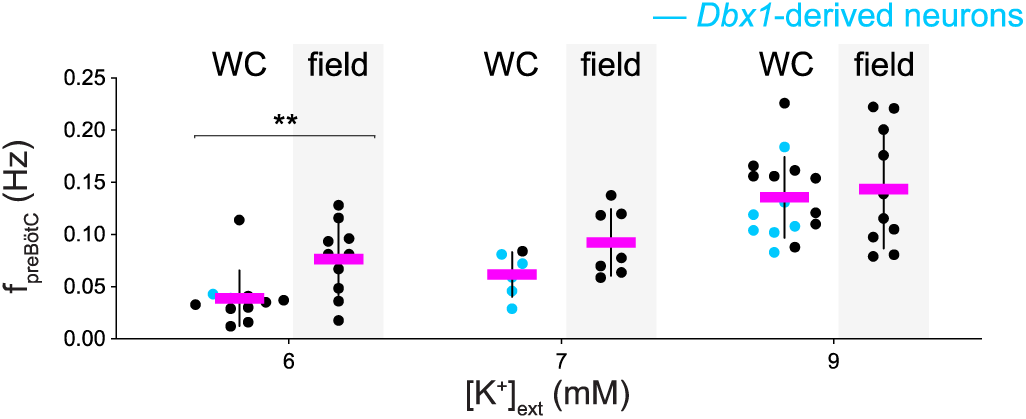
Comparison of frequency of preBötC composite rhythm measured intracellularly and via field recordings. Frequency of preBötC composite rhythm is measured intracellularly and via field recordings at three levels of external K^+^ concentration, i.e., [K^+^]_o_: 6, 7, and 9 mM. At each [K^+^]_o_ level, whole-cell (WC) is at left and the preBötC field recording is at right (with light gray background shading). Cyan symbols represent measurements from *Dbx1*-derived preBötC neurons in Dbx1;Ai148 reporter mice. Asterisks signify statistical significance at p < 0.05.

### Burstlets occur in subset s of inspiratory preBötC neurons

To investigate how many and which neurons participate in burstlets, we recorded inspiratory *Dbx1*-derived preBötC neurons in Dbx1;Ai148 slices while simultaneously monitoring XII motor output (Figs. 7 and 8). We recorded 3-9 imaging planes per preBötC with 12 ± 7 active neurons per plane (range 3-27) for an average of 62 ± 20 inspiratory neurons recorded per Dbx1;Ai148 slice. Then, we manipulated [K^+^]_o_ to examine burst (9 mM; n = 6 slices) and burstlet (7 mM, n = 6 slices; 6 mM, n = 2 slices) rhythms.

**Figure 7.**
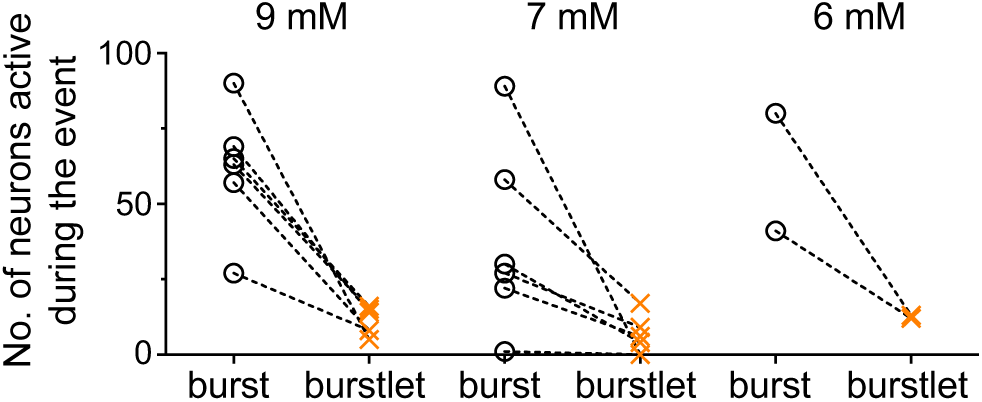
Burstlets occur in subset of preBötC inspiratory neurons. Group data showing the number of neurons active during bursts and burstlets from Dbx1;Ai148 slices at 9, 7, and 6 mM external K^+^ concentration, i.e., [K^+^]_o_. Active neuron counts are illustrated for each slice preparation for bursts (open circles) and burstlets (orange X symbols). Only two slices were sufficiently rhythmically active at 6 mM [K^+^]_o_ to obtain reliable measurements of burstlets within the 2-min recording duration of imaging time series. Insets in A and B show select burstlets.

**Figure 8.**
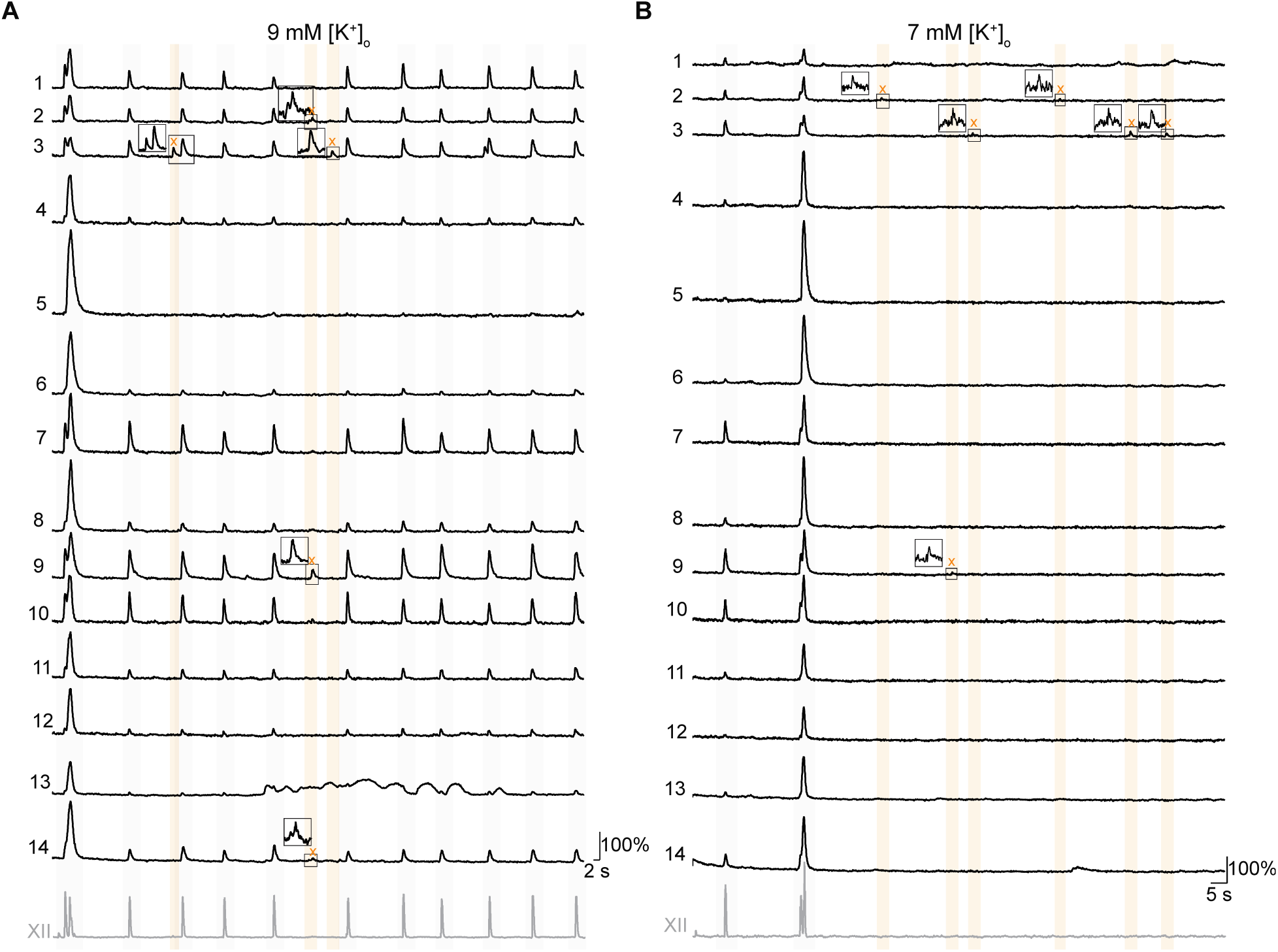
Burstlets occur in a dynamic subset of preBötC inspiratory neurons. Bursts and burstlets in a group of 14 *Dbx1*-derived preBötC neurons in a typical Dbx1;Ai148 mouse slice preparation with inspiratory XII output (lowest traces, gray). Inspiratory cycles are background shaded in light gray. Burstlet cycles are background shaded in light orange. A and B show population activity in 9 and 7 mM external K^+^ concentration, i.e., [K^+^]_o_, respectively.

At 9 mM [K^+^]_o_, 20% ± 9% of the *Dbx1*-derived inspiratory neurons were active during burstlets (Fig. 7). Additionally, a neuron that was active during one burstlet was not always active during other burstlets (Fig. 8A, neurons 2, 3, 9, 14). Upon changing to 7 mM [K^+^]_o_, 17% ± 15% of the *Dbx1*-derived inspiratory neurons were active during burstlets (Fig. 7). Again, a neuron that participated in one burstlet did not always participate in the next burstlet (Fig. 8B, neurons 2, 3, 9). Upon changing to 6 mM [K^+^]_o_, 23% of *Dbx1*-derived neurons were active during burstlets (Fig. 7). These data show that the subset of *Dbx1* preBötC neurons that participates in burstlets constitutes 17-23% of the population active during inspiratory bursts and that the composition of the burstlet-active subset varies from cycle to cycle.

## DISCUSSION

Inspiratory breathing movements emanate from neural activity in the preBötC but the cellular and synaptic mechanisms underlying its rhythmicity remain incompletely understood and – some might argue – misunderstood. Specifically, there appears to be a dichotomy between the mechanisms underlying rhythmogenesis and those mechanisms governing motor output pattern. Here we investigate this rhythm-pattern dichotomy to unravel the neural mechanisms of preBötC functionality.

### Defunct theories of inspiratory rhythmogenesis and the viability of burstlets to explain it

Theories of rhythmogenesis fall into three camps. The first posits a ring of mutually inhibitory neurons that generates sequential phases of the breathing cycle including inspiratory bursts in the preBötC (Ausborn et al. 2018; Richter 1982; J. C. Smith et al. 2007; Jeffrey C Smith et al. 2013). The second theoretical framework emphasizes *pacemaker* neurons whose voltage-dependent conductances enable rhythmic bursting; the synchronization of pacemakers presumably serves as a template for network activity (Butera, Rinzel, and Smith 1999a, 1999b; Feldman and Cleland 1982; S. M. Johnson et al. 1994; Ramirez, Tryba, and Peña 2004; Jens C. Rekling and Feldman 1998). The third theory, dubbed a *group pacemaker*, posits that recurrent synaptic activity triggers mixed-cationic conductances to produce inspiratory bursts (Del Negro and Hayes 2008; Jens C. Rekling and Feldman 1998; J. C. Rekling, Champagnat, and Denavit-Saubie 1996; Rubin et al. 2009). However, disinhibition of the preBötC (Baertsch, Baertsch, and Ramirez 2018; Brockhaus and Ballanyi 1998; Cregg et al. 2017; Janczewski et al. 2013; Marchenko et al. 2016; Shao and Feldman 1997; Sherman et al. 2015), as well as attenuation of pacemaker conductances (Del Negro et al. 2002, 2005; Koizumi and Smith 2008; Pace et al. 2007; Peña et al. 2004) and mixed-cationic conductances (Koizumi et al. 2018; Picardo et al. 2019) neither perturbs the frequency in the predicted manner nor stops breathing *in vivo* or inspiratory rhythms *in vitro*, which therefore falsifies all three rhythmogenic mechanisms. Nevertheless, the key to understanding rhythmogenesis may be found in what these theories get wrong: inextricable neural bursts that culminate the inspiratory phase of the cycle.

To consider the iconoclastic notion of rhythmogenesis in the absence of bursts we, like Kam & Feldman (2013a, 2015), focus on the preinspiratory phase that ordinarily leads inexorably to bursts and motor output. The preinspiratory phase is a hallmark of rhythmogenesis, putatively marking early-activating rhythmogenic respiratory interneurons (Carroll and Ramirez 2012; Carroll, Viemari, and Ramirez 2012; Onimaru, Arata, and Homma 1987, 1988; J. C. Rekling, Champagnat, and Denavit-Saubie 1996; J. C. Smith et al. 1990). Concurrent excitation of 4-9 excitatory preBötC interneurons *in vitro*, by photolytic glutamate uncaging (Kam, Worrell, Ventalon, et al. 2013; J. C. Rekling, Champagnat, and Denavit-Saubie 1996; J. C. Smith et al. 1990; Sun et al. 2019), can effectively trigger a preBötC network burst after a latency of 100-400 ms, similar to the duration of preinspiratory activity, shown in our Fig. 5 and by others. There are two important take-aways: first, small numbers (<10) of coactive neurons can trigger a burst; second, the burst occurs after sufficient time for percolation of network interactions to reach threshold. Kam and Feldman (2013a) divorced preinspiratory activity from bursts, showing that rhythmic burstlets remained in their absence, and they argued that burstlets represent the rhythmogenic substrate.

We also observed spontaneous preBötC field activity like burstlets that do not lead to XII output at all integer levels between 3 and 9 mM [K^+^]_o_. Further, by manipulating the excitability in the preBötC, we could detach the preinspiratory and inspiratory components of preBötC burst, as well as affect their prevalence. Field and whole-cell recordings from preBötC neurons showed that the rise time, duration, and amplitude of burstlets matches preinspiratory activity. These data agree with the burstlet hypothesis that both burstlet and preinspiratory activity indeed follow a common rhythmogenic mechanism.

### How many constituent neurons activate during bursts and burstlets?

Kam and Feldman used loose-patch recordings to show that ∼89% of preBötC neurons active during bursts also participate in burstlets. Here, using photonics to monitor an average of 62 *Dbx1*-derived inspiratory burst-active neurons, we found ∼20% participate in burstlets. Additionally, we noted disparity between the frequency of preBötC composite rhythm between field recordings and whole-cell recordings at 6 mM [K^+^]_o_. Whereas field recordings reflect activity among many preBötC neurons, whole-cell recordings focus on one constituent preBötC neuron. As excitability decreases, fewer neurons participate in burstlets but field recordings still detect them. In contrast, any neuron singled out for whole-cell recording is less likely to be part of the burstlet-active subpopulation. Thus, one finds higher frequency of composite rhythm, compared to whole-cell recordings at an intermediate level of excitability (i.e., 6 mM). Our data indicate that the subset of burstlet-active neurons is inconstant and lower than 89%. We have estimated the size of the rhythmogenic population to be 560-650 preBötC neurons (Hayes, Wang, and Del Negro 2012; Wang et al. 2014) so the burstlet-active subpopulation numbers between 112-130 (20%) and 500-580 (89%). That seemingly large range can explain why burstlet amplitude is voltage-dependent: increasing excitability can recruit potentially hundreds of additional constituent neurons to the burstlet-active subpopulation in the preBötC. Whether or not the fraction of burstlet-active preBötC neurons is closer to 20% or 89%, very few (<10) coactive preBötC neurons can trigger full bursts and motor output (Kam, Worrell, Ventalon, et al. 2013; Sun et al. 2019) so the relative fraction of burstlet-active neurons may not be a critical parameter governing network activity.

### Burstlet mechanisms

Here, the frequency of preBötC composite rhythm was voltage dependent. In contrast, the initial burstlet report (Kam et al. 2013a) showed no statistically significant disparity between the frequency of preBötC field activity at 6 vs. 9 mM [K^+^]_o_, yet there was a disparity for 3 vs. either 6 or 9 mM. This left the question open as to whether burstlet rhythm might be some form of synchronized voltage-independent biochemical oscillator in constituent neurons. Monotonically increasing frequency of preBötC composite rhythm as a function of [K^+^]_o_ now rules out that possibility.

So what mechanism does give rise to burstlets? Our whole-cell recordings show temporal summation of EPSPs during burstlets. We held membrane potential at –60 mV, which imposes steady-state deactivation of the persistent Na^+^ current (Del Negro et al. 2002; Ptak et al. 2005; Yamanishi et al. 2018). Therefore, burstlets do not reflect voltage-dependent bursting properties and do appear to reflect recurrent synaptic excitation.

In general, network oscillators (distinct from pacemaker or inhibition-based models) rely on recurrent excitation among constituent rhythmogenic neurons (Grillner 2006). Modifying the neuronal excitability levels via [K^+^]_o_ influences the relative fraction of spontaneously active neurons in the network, and, for silent neurons, the proximity of baseline membrane potential to spike threshold. Increasing excitability therefore magnifies the number of neurons interacting, facilitates summation of their synaptic drives, and accelerates the process of recurrent excitation to directly influence frequency. We conclude that burstlets are not only rhythmogenic, but also follow dynamics of recurrent excitation, i.e., a network oscillator model of rhythmogenesis.

### Pattern generation

Burstlet amplitude is voltage dependent too. Above, we inferred that additional constituent neurons (perhaps ∼100s) are recruited as excitability increases. In contrast, the amplitude of preBötC bursts and XII motor output are not voltage dependent across [K^+^]_o_ levels. We conclude that preBötC activity, once passing threshold, triggers a cascade that activates all (or nearly all) constituent preBötC neurons, and also activates premotor and motor neurons. That cascade probably depends on synaptic connectivity among pattern-related preBötC neurons and premotor neurons outside of the preBötC, but not preBötC excitability *per se*.

We cannot yet specify how activity during the preinspiratory phase reaches a threshold for burst generation. It may have to do with a quorum: a certain number of rhythmogenic interneurons must be active (i.e., spiking) to trigger an irreversible cascade that ostensibly activates all (or nearly all) preBötC neurons. Or, it may have to do with synchrony: a certain number of rhythmogenic interneurons must be spiking in sync to trigger that cascade. The first scenario focuses on mass action of constituent interneurons wherein temporal precision is inconsequential. The second scenario focuses on phasic precision rather than mass action. Our present data cannot distinguish which mechanism is at work but Ashhad and Feldman argue for the preeminent importance of synchrony in burst generation (Ashhad and Feldman 2019). Given that burstlet amplitude is voltage-dependent and the number of constituent neurons participating in burstlets may vary (∼100s) at any given level of excitability, the quorum model seems less feasible.

There is an existing framework for understanding both rhythm and pattern generation of the preBötC. *Dbx1*-derived preBötC neurons are inspiratory rhythmogenic (Baertsch, Baertsch, and Ramirez 2018; Bouvier et al. 2010; Cui et al. 2016; Gray et al. 2010; Koizumi et al. 2016; Vann et al. 2016, 2018; Wang et al. 2014), playing key preinspiratory and burst-generating roles (Cui et al. 2016; Picardo et al. 2013) and some serving exclusively premotor function (Revill et al. 2015). It may be possible to identify subsets of predominantly rhythmogenic versus predominantly premotor or pattern-related *Dbx1*-derived preBötC neurons based on intrinsic membrane properties (Picardo et al. 2013) or neuropeptide somatostatin expression (Cui et al. 2016). Those two classification schemes are not mutually exclusive because many somatostatin- and somatostatin receptor-expressing preBötC neurons are *Dbx1*-derived (Gray et al. 2010).

It is also possible that particular ion channels serve in a pattern-related capacity within core rhythmogenic neurons. We and others recently characterized transient receptor potential (Trp) channels in *Dbx1*-derived preBötC neurons whose activation amplifies inspiratory burst magnitude (Koizumi et al. 2018; Picardo et al. 2019) and we exclusively demonstrated that Trp channels in the preBötC maintain the tidal volume of inspiratory breaths in unanesthetized adult mice (Picardo et al. 2019).

Our results affirm the ideas presented in the burstlet hypothesis (Kam et al., 2013a) that rhythm and pattern generation are discrete processes, which nevertheless both begin in the preBötC. Burstlets, subthreshold from the standpoint of motor discharge, appear to reflect the core rhythmogenic mechanism involving recurrent synaptic excitation.

